# Sex-Dependent Proteomic Remodelling During *ex vivo* Degeneration of Young and Aged Murine Peripheral Nerves

**DOI:** 10.64898/2026.08.04.742743

**Authors:** Dario Lucas Bergmann, Emilio Cirri, Joanna M. Kirkpatrick, Erika Kelmer Sacramento, Leonie Karoline Stabenow, Nova Oraha, Leopold Böhm, Martin Walter, Reinhard Bauer, Helen Morrison

## Abstract

**Introduction:** Peripheral nerve ageing leads to profound proteomic remodelling, with shifts in metabolic and inflammatory signalling pathways resembling changes that occur during nerve degeneration and regeneration following injury. Moreover, aged nerves exhibit impaired degeneration and regeneration, contributing to age-related peripheral neuropathies that show sex-specific differences in prevalence. However, it remains unclear whether these alterations arise from intrinsic nerve changes or an altered systemic environment. Therefore, we investigated the impact of sex on age-related proteome changes and nerve-intrinsic proteomic responses in young and aged male and female nerves using an *ex vivo* degeneration model.

**Methods:** Mass spectrometry-based proteomics were performed on young and old nerves from male and female animals, as well as on contralateral nerves after seven days of *ex vivo* nerve degeneration. A comparative bioinformatic analysis was then used to identify changes during ageing and *ex vivo* nerve degeneration that were independent of sex, as well as changes that were sex-specific.

**Results:** *Ex vivo* nerve degeneration induced extensive proteome remodelling in mouse sciatic nerves that was largely independent of age and sex. Principal component and clustering analyses clearly separated intact from degenerated nerves, while revealing only subtle age- and sex-related effects, with more pronounced ageing-associated changes in males. Approximately 20% of age-regulated proteins and 7–10% of degeneration-regulated proteins exhibited sex-specific expression patterns. Degeneration was characterised by increased abundance of lysosomal and repair-associated proteins alongside reduced myelin and axonal proteins, consistent with active tissue remodelling. In aged nerves, impaired protein clearance and partial pre-activation of degeneration-associated pathways suggested altered injury responses.

Comparative analyses demonstrated positive correlations of protein abundance changes between *ex vivo* and *in vivo* degeneration datasets, although the temporal dynamics were altered in aged nerves. Pathway enrichment analyses identified coordinated regulation of metabolic, RNA-processing and vesicular transport pathways, while ageing was associated with enhanced immune signalling and reduced lipid metabolism. Sex-specific analyses revealed stronger inflammatory signatures in males, whereas females exhibited enrichment of metabolic pathways, including folate biosynthesis.

**Conclusion:** These findings reveal distinct sex-specific molecular features of peripheral nerve ageing, characterised by enhanced inflammatory signalling in males and metabolic adaptations that may confer resilience in females. Our datasets provide a comprehensive molecular resource of sex-dependent changes in peripheral nerve ageing and nerve-intrinsic injury responses, offering a foundation for identifying therapeutic strategies to promote healthy peripheral nerve ageing.

**Plain English summary:** Age-related peripheral neuropathies are common disorders that can cause pain, numbness, weakness and reduced mobility, affecting millions of people worldwide. They become more common from around the age of 50 and affect men and women differently. These conditions are thought to result from age-related changes in the structure and function of peripheral nerves, which reduce their ability to repair themselves after injury. In this study, we used advanced protein analysis (proteomics) to investigate how ageing affects peripheral nerves in male and female mice. We also used an *ex vivo* model, in which nerves are studied outside the body, to examine how age and sex influence the molecular changes that occur during nerve degeneration. We found that degeneration caused widespread changes in the proteins present in the sciatic nerve in both young and old mice. Most of these changes were similar in males and females, but some important differences emerged. Male nerves showed stronger signs of inflammation, whereas female nerves showed increased activity of metabolic pathways, including those involved in folate metabolism. Ageing nerves also appeared less able to remove damaged material and showed signs of activating degeneration-related processes even before injury. Overall, the *ex vivo* model reproduced many of the molecular changes seen after nerve injury in living animals, although it did not fully capture the inflammatory response, suggesting that signals from the rest of the body, including factors carried in the blood, also contribute to nerve degeneration.

**Highlights:**

- *Ex vivo* nerve degeneration caused major protein changes in young and old mouse sciatic nerves.
- Most degeneration-related protein changes were shared between males and females.
- Ageing altered the nerve proteome, with stronger ageing-related shifts in males.
- Male nerves showed stronger inflammatory and immune-related signatures.
- Female nerves showed enrichment of metabolic pathways, including folate biosynthesis, and *ex vivo* degeneration did not fully reproduce the inflammatory response seen after injury *in vivo*.

## Background

Ageing of the peripheral nervous system (PNS) leads to morphological, molecular and functional changes that ultimately affect the homeostasis and regenerative potential of the PNS, which results in the development of age-associated peripheral neuropathies (1–5). However, neuronal growth and axonal sprouting remain unaffected by ageing (5, 6), suggesting that impaired regeneration results primarily from age-related dysfunction of non-neuronal cells and alterations to the nerve microenvironment. Previous studies have shown that **both** Schwann cells and macrophages exhibit impaired myelin debris clearance following PNS injury (3, 5, 7), a process that is essential for successful nerve regeneration (8, 9). While macrophages primarily rely on receptor-mediated phagocytosis (RMP), Schwann cells use both RMP and the specialised autophagic process “myelinophagy” (7, 8, 10). Cell culture studies with isolated murine Schwann cells have demonstrated reduced myelinophagic capacity in aged cells (5). However, in a heterochronic parabiosis model, myelin debris clearance improved when an old mouse shared the blood circulation of a young mouse, suggesting that systemic factors can partially restore regenerative processes. Although these findings were obtained in the central nervous system (11), they highlight the potential contribution of the systemic environment to age-related deficits in nerve repair.

Mass spectrometry-based proteomics provides a powerful approach to quantify the molecular dynamics of myelin degeneration and degradation following peripheral nerve injury *in vivo* (12). Furthermore, nerve crush injury leads to profound proteomic remodelling of multiple signalling networks, with time-point-specific proteome profiles (13). Consequently, proteomic analysis of peripheral nerve degeneration in an *ex vivo* model offers the opportunity to distinguish nerve-intrinsic age-related proteomic changes from those driven by the aged systemic environment.

Moreover, although sex-specific changes in ‘inflammageing’ have been extensively studied, with women generally exhibiting **lower age-related increases in inflammatory biomarkers**, potentially contributing to their reduced risk of cardiovascular disease and longer life expectancy (14), whether similar sex-specific molecular changes occur in the ageing peripheral nervous system to our acknowledge remains unknown. Female rats exhibit faster peripheral nerve regeneration than males (15), possibly reflecting hormonal influences. For example, 17β-estradiol enhances Schwann cell differentiation, remyelination and regeneration after sciatic nerve crush injury in mice (16). In humans, women also appear to be more resistant to denervation following neural injury, showing better functional recovery in patients with facial paralysis (17). However, evidence from mouse models is less consistent. Although one study found no sex differences in axonal degeneration or regeneration three days after sciatic nerve crush injury, transcriptomic analysis revealed numerous sex-dependent differences in genes associated with nerve regeneration (18). In addition, Zhou et al. demonstrated that, following sciatic nerve injury, inflammatory proteins were systemically upregulated in the blood, with a more modest increase in female mice, suggesting that sex-specific inflammatory responses may contribute to differences in neuropathic pain development (19).

Previously, we demonstrated that ageing induces proteomic changes in peripheral nerves that resemble those observed following injury, an effect that was partially attenuated by long-term caloric restriction (13). Taken all together these findings led us to hypothesize that the intrinsic molecular response to nerve injury differs between young and aged nerves. In addition, despite increasing recognition of sex differences in inflammation and nerve regeneration, sex-specific molecular changes during peripheral nerve ageing have not been comprehensively characterized. Therefore, we extended our previous proteomics studies by combining an *ex vivo* nerve degeneration model with quantitative proteomics to investigate how ageing and sex influence nerve-intrinsic proteomic remodeling during peripheral nerve degeneration.

## Materials and Methods

### Experimental animals

All animals were on a C57BL/6 J background. Mice were group housed, with free access to standard chow and water and maintained with 12 h light and dark cycles. Temperature was 20 ± 2 °C during the experimental period. The animal procedures were performed according to the guidelines from Directive 2010/63/EU of the European Parliament on the protection of animals used for scientific purposes and tissue harvesting from mice was performed according to § 11 TierSchG (Haltungserlaubnis) at the FLI. Mice were considered to be old/geriatric at ∼120 weeks and considered young at around 8-12 weeks.

### *Ex vivo* nerve degeneration assay

This experiment was performed as previously described (8, 20). In brief, sciatic nerves from 6-months old, adult mice were removed, cleared from superfluous connective tissue, and cut into two halves before placing into DMEM+5% FBS+ 2% P/S. The nerves were allowed to degenerate for 7 days before they were snap-frozen in liquid nitrogen or fixed in 4% ice-cold PFA. Intact sciatic nerves from the contralateral side were used as control tissue from indicated genotypes and were directly snap-frozen or fixed after harvesting.

### Protein isolation from sciatic nerve tissue

Snap frozen intact or degenerated nerves were thawed on ice and were homogenized in a Precellys® 24 homogenizer (Bertin Instruments, Montignyle-Bretonneux, France) in Pierce RIPA buffer (Thermo Fisher Scientific Inc., Waltham, MA, USA) before further processing for proteomic analysis.

### Proteomics of sciatic nerves

#### Sample preparation and LC-MS/MS data independent acquisition/analysis (DIA)

For proteomics analysis, samples were sonicated (Bioruptor Plus, Diagenode, Belgium) for 10 cycles (30 sec ON/60 sec OFF) at high setting, at 20 °C, followed by boiling at 95°C for 5 min. Reduction was followed by alkylation with iodoacetamide (IAA, final concentration 15 mM) for 30 min at room temperature in the dark. Protein amounts were estimated following an SDS-PAGE gel of 10 µL of each sample against an in-house cell lysate of known quantity. 30 µg of each sample was taken along for digestion. Proteins were precipitated overnight at −20 °C after addition of 8x volume of ice-cold acetone. The following day, the samples were centrifuged at 20800x g for 30 min at 4 °C and the supernatant carefully removed (Eppendorf 5810R, Eppendorf AG, Germany). Pellets were washed twice with 300 µL ice-cold 80% (v/v) acetone in water then centrifuged at 20800x g at 4 °C for 10 min. After removing the acetone, pellet were air-dried before addition of 25 µL of digestion buffer (1M Guanidine, 100 mM HEPES, pH 8). Samples were resuspended with sonication as explained above, then LysC (Wako) was added at 1:100 (w/w) enzyme:protein ratio and digestion proceeded for 4 h at 37 °C under shaking (1000 rpm for 1 h, then 650 rpm). Samples were then diluted 1:1 with MilliQ water and trypsin (Promega) added at at 1:100 (w/w) enzyme:protein ratio. Samples were further digested overnight at 37 °C under shaking (650 rpm). The day after, digests were acidified by the addition of TFA to a final concentration of 10% (v/v), heated at 37 °C and then desalted with Waters Oasis® HLB µElution Plate 30 µm (Waters Corporation, MA, USA) under a soft vacuum following the manufacturer instruction. Briefly, the columns were conditioned with 3×100 µL solvent B (80% (v/v) acetonitrile; 0.05% (v/v) formic acid) and equilibrated with 3×100 µL solvent A (0.05% (v/v) formic acid in Milli-Q water). The samples were loaded, washed 3 times with 100 µL solvent A, and then eluted into 0.2 mL PCR tubes with 50 µL solvent B. The eluates were dried down using a speed vacuum centrifuge (Eppendorf Concentrator Plus, Eppendorf AG, Germany). Dried samples were stored at −20°C until analysis.

Prior to analysis, samples were reconstituted in in MS Buffer (5% acetonitrile, 95% Milli-Q water, with 0.1% formic acid) and spiked with iRT peptides (Biognosys, Switzerland). Peptides were separated in trap/elute mode using the nanoAcquity MClass Ultra-High Performance Liquid Chromatography system (Waters, Waters Corporation, Milford, MA,USA) equipped with a trapping (nanoAcquity Symmetry C18, 5 μm, 180 μm × 20 mm) and an analytical column (nanoAcquity BEH C18, 1.7 μm, 75 μm × 250 mm). Solvent A was water and 0.1% formic acid, and solvent B was acetonitrile and 0.1% formic acid. 1 µl of the sample (∼1 μg on column) were loaded with a constant flow of solvent A at 5 μl/min onto the trapping column. Trapping time was 6 min. Peptides were eluted via the analytical column with a constant flow of 0.3 μl/min. During the elution, the percentage of solvent B increased in a nonlinear fashion from 0–40% in 120 min. Total run time was 145 min. including equilibration and conditioning. The LC was coupled to either to an Orbitrap Q-Exactive HFX (Thermo Fisher Scientific, Bremen, Germany) using the Proxeon nanospray source. The peptides were introduced into the mass spectrometer via a Pico-Tip Emitter 360-μm outer diameter × 20-μm inner diameter, 10-μm tip (New Objective) heated at 300 °C, and a spray voltage of 2.2 kV was applied. The capillary temperature was set at 300°C. The radio frequency ion funnel was set to 30%.

For data independent acquisition (DIA), full scan mass spectrometry (MS) spectra with mass range 350–1650 m/z were acquired in profile mode in the Orbitrap with resolution of 120,000 FWHM. The default charge state was set to 3+. The filling time was set at maximum of 60 ms with limitation of 3 × 106 ions. DIA scans were acquired with 40 mass window segments of differing widths across the MS1 mass range. Higher collisional dissociation fragmentation (stepped normalized collision energy; 25, 27.5, and 30%) was applied and MS/MS spectra were acquired with a resolution of 30,000 FWHM with a fixed first mass of 200 m/z after accumulation of 3 × 106 ions or after filling time of 35 ms (whichever occurred first). All data were acquired in profile mode. For data acquisition and processing of the raw data Xcalibur 4.0 (Thermo) and Tune version 2.9 were used.

### Proteomic data processing

Acquired data were processed using Spectronaut Professional v13.10 (Biognosys AG). DIA raw data were analyzed using the directDIA pipeline in Spectronaut v.20 (Biognosys, Switzerland) with BGS settings besides the following parameters: Proteotypicity Filter = Only Protein Group Specific; Major Group Quantity = Median peptide quantity; Major Group Top N = OFF; Minor Group Quantity = Median precursor quantity; Minor Group Top N = OFF; Data Filtering = Qvalue sparse; Normalization Strategy = Local normalization; Row Selection = Automatic. The data were searched against a species specific (Mus musculus, 16,747 entries, v. 160106) and a contaminants (247 entries) Swissprot database. Relative protein quantification was performed in Spectronaut using a pairwise t-test performed at the precursor level followed by multiple testing correction according to Benjamini-Hochberg. The data (candidate table) and data reports (protein quantities) were then exported for further data analyses. The obtained dataset has been deposited on the MassIVE repository (MSV000101693).

### Bioinformatic data analysis

All data analysis steps were completed with R(21). Data wrangling steps were performed using primarily dplyr functions from the tidyverse package(22). PCA and further clustering analysis was performed on log_2_ transformed PG.Quants using the prcomp function and visualization was performed using the factoextra package for proteomics data. Differentially expressed proteins were obtained by using a |log2FC| ≥ 0.58 and q ≤ 0.05 cutoff based on Spectronaut analyses as described before. The following experimental comparisons of interest were defined for the dataset: A1 = Young degenerated male / Young intact male, A2 = Young degenerated female /

Young intact female, B1 = Old degenerated male / Old intact male, B2 = Old degenerated female / Old intact female, C1 = Old intact male / Young intact male, C2 = Old intact female / Young intact female, D1 = Young intact female / Young intact male, D2 = Old intact female / Old intact male and labelled as indicated in the respective figure legends. Weighted gene/protein correlation network analysis (WGCNA) was performed using the WGCNA(23) and CEMiTool(24) R packages. All proteins were included in the WGCNA analysis. First, clustering of module eigengenes and visualization with a dendrogram was performed to identify a biologically reasonable threshold, for both the transcriptome and the proteome a threshold of 0.8 eigengene correlation was used. Subsequently, signed networks were constructed for better biological interpretability and biweight midcorrelation was used as a robust correlation measure. The final WGCNA function with previously mentioned settings was in the end run using the cemitool() function that chooses the soft-thresholding power β automatically to achieve scale-free topology(24). Gene set enrichment analysis (GSEA) was performed with the “fgsea” package(25). Heatmaps were created using the “ComplexHeatmap” package(26) of row-wise z-score normalized values from the log_2_ PG.Quants. The resulting dendrogram was then resorted with “dendsort” package(27). Gene ontology and KEGG pathway enrichment analyses were conducted using the clusterProfiler R package(28) based on an over-representation analysis (ORA). GO-network function visualization using an enrichment map was done using the clusterProfiler R package(28). Visualization of significantly enriched GO-terms (q < 0.1) and clustering was done using the “simplifyEnrichment” package(29) with kmeans clustering as method for semantic similarity clustering. All results from enrichment analyses are provided in **Supplementary file 2**. Correlation network using UniProt IDs was computed using the correlate()(function from the “corrr” package and the network was visualised by creating an igraph object(30) and then using the ggraph package for visualization. Interferon network analysis was performed as previously described(31) using the network clusters identified by Mostafavi et al.(32). The myelin protein set was retrieved from(12) and proteins annotated to GO-terms axon and lysosome were retrieved using the biomaRt package(33). All z-scored protein intensities as well as files with the results from the differential expression analysis for all datasets are provided in **Supplementary file 1**.

### Use of artificial intelligence (LLMs) for manuscript preparation

During the preparation of this work, the authors used OpenEvidence, ChatGPT and DeepL for literature search (OE and CG) and to improve language, conciseness and readability of the content (CG and DL). After using these tools, the authors reviewed and edited the content as needed and take full responsibility for the content of the publication.

### Statistical procedures

All analyses were performed using Graphpad Prism 8.4 or R(21). Shapiro-Wilk normality test was carried out before further statistical analysis was performed. For analysis of changes in myelin, axonal and lysosomal proteins, Kruskal-Wallis tests were performed with subsequent post-hoc tests with FDR correction of the p-values using the two-stage linear step-up procedure of Benjamini, Krieger and Yekutieli. Statistical tests for proteomic analysis and functional annotation and enrichment analyses were performed as indicated in the respective material and methods section. Statistical significance was accepted if p ≤ 0.05, unless otherwise stated in the respective figure legend.

## Results

### *Ex vivo* nerve degeneration leads to profound proteome remodelling in young and old mouse nerves

To characterise global proteomic changes during peripheral nerve ageing and degeneration (**Figure 1A**), as well as differences between sexes, we performed a principal component analysis (**Figure 1B**). Along the first principal component, which explains 52.1% of the total variance, a clear separation is visible between degenerated and intact nerves for both ages and sexes, highlighting the profound impact of *ex vivo* nerve degeneration on the sciatic nerve proteome. Furthermore, old nerves are separated from young nerves along the second principal component. Though this separation remains after *ex vivo* degeneration, young and old degenerated nerves are closer to each other than to their intact counterparts. Additionally, old male intact nerves are separated from old female nerves along PC2, suggesting potentially more advanced ageing in males compared to females. To assess sample quality, we plotted the log₂ protein quantities for all samples and performed a correlation analysis of all samples with the proteomes of all other individual samples (**Figures 1C and 1D**). All samples showed a uniform distribution (**Figure 1D**), and all degenerated nerves exhibited high correlations with each other, while demonstrating only moderate to low correlations with intact nerves. This indicates the high reproducibility of *ex vivo* nerve degeneration and the good quality of the samples. Furthermore, hierarchical clustering of the correlations showed a separation between old and young nerves in both intact and degenerated nerves, highlighting the effect of ageing on the sciatic nerve proteome (**Figure 1C**). We therefore performed a weighted gene co-expression/correlation network analysis to identify clusters of proteins that were highly correlated. This analysis identified three clusters: two that were either upregulated or downregulated during *ex vivo* nerve degeneration (Cluster M1 and cluster M2, respectively), and one that contained proteins that were upregulated in old nerves and unaffected by degeneration (Cluster M3; **Figure S1A**). Enrichment analyses of the co-expression modules across all experimental groups showed slightly stronger up- and downregulation of the injury-regulated modules in male mice compared to female mice, as well as slightly weaker regulation of the injury-regulated modules in old mice compared to young mice (**Figure S1B**). Additionally, the module affected solely by ageing was slightly more enriched in old male mice than in old female mice, indicating stronger ageing-induced changes in males than in females (**Figure S1B**).

**Fig. 1.**
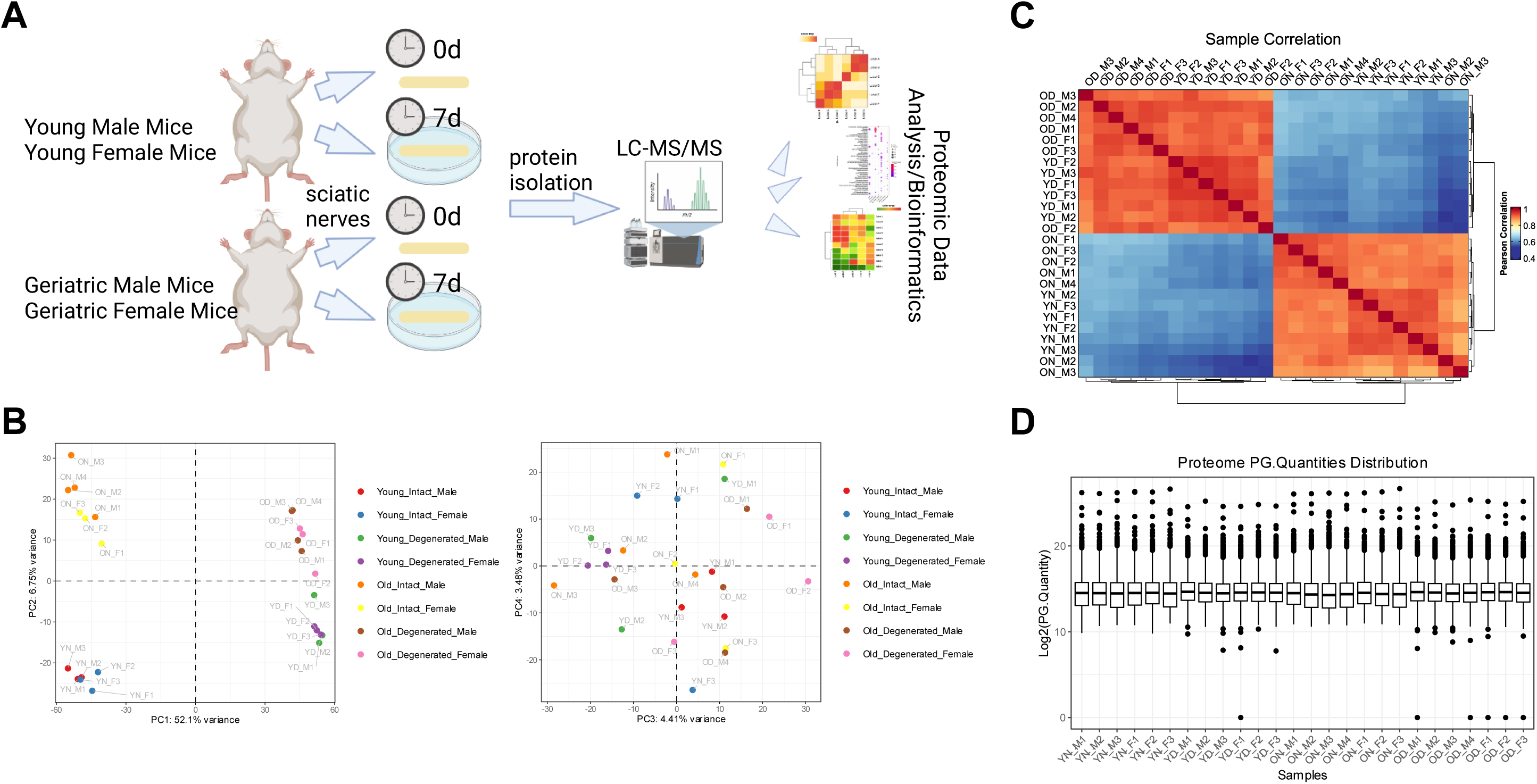
Principal component analysis (PCA), sample correlation and proteome data distribution. **A** Schematic experimental workflow for *ex vivo* nerve degeneration assay. **B** Principal component analysis (PCA) of intact and *ex vivo* degenerated nerves (n = 3 per group, except old male mice n = 4). In the left plot PC1 and PC2 are depicted, in the right plot PC3 and PC4 are depicted. **C** Sample correlation heatmap using Pearson correlation coefficients. **D** Boxplot of the log_2_ Protein Group Quantities (PG.Quantities) across all samples.

### Characterisation of the degenerative response during *ex vivo* nerve degeneration, compared with *in vivo* nerve degeneration

To investigate whether proteomic changes with *ex vivo* degeneration differ between male and female nerves, we performed a correlation analysis of the log₂ fold changes of degenerated male and female nerves relative to intact male and female nerves in each age group, and calculated Spearman’s ρ correlation coefficients. Strong correlations were observed for both age groups, highlighting that the general proteomic response to *ex vivo* nerve degeneration remains consistent throughout the ageing process (**Figure 2A**). However, around 7–10% of proteins were uniquely regulated in either male or female nerves, with male nerves showing slightly more unique changes (**Figure 2A**). In addition, ageing-induced changes in intact nerves showed a moderate positive correlation between males and females, despite considerable sex-specific changes (18–23% of all proteins) (**Figure 2A**). Among the proteins that were upregulated with degeneration, independent of age or sex, were lysosomal proteins (*CD68* and *Lamp1*) and *Arg1*, a marker protein of activated macrophages (Figure 2A). Conversely, several inflammatory proteins, such as complement proteins (*C3*) and immunoglobulins, were downregulated in degenerated nerves. Furthermore, we identified a classical regulator of Schwann cell repair cell phenotype, *Shh*, to be strongly induced in young nerves with degeneration, but only weakly in old nerves with degeneration, while protein levels were already upregulated in intact, old nerves in comparison to young nerves and, in addition, in old female nerves stronger than in old male nerves (**Figure 2A**). To assess signs of neurodegeneration and myelin degradation, we examined how myelin, axonal and lysosomal proteins are regulated in all experimental groups. In general, we identified a decrease in myelin protein levels alongside an increase in lysosomal proteins and degradation, with no differences between sexes (**Figure 2B**). However, myelin protein levels decreased significantly in old nerves to the same extent as in young nerves during degeneration. There was also only a trend towards a decrease in myelin protein levels with degradation in old, male nerves, which indicates potentially defective clearance mechanisms in the old, male PNS (**Figure 2B**). Axonal proteins were also strongly downregulated during degeneration and to some extent during ageing, but only in male mice, while axonal proteins were preserved during ageing in female mice (**Figure 2B**). Furthermore, we investigated the overlap between proteome profiles during *ex vivo* and *in vivo* nerve degeneration by calculating correlation coefficients of all comparisons reported in this study to previously reported post-injury datasets at different time points after injury (3dpc, 7dpc and 28dpc) (13) and constructing a correlation network (**Figure 2C, Supplementary table 1**). We observed a structured organisation in which post-injury proteome profiles positively correlated with each other for both *in vivo* and *ex vivo* degeneration. This suggests that similar degeneration-induced changes occur during *ex vivo* degeneration compared to *in vivo* degeneration. Furthermore, the correlation coefficients of the *ex vivo* degeneration datasets were higher for the *in vivo* 7 dpc dataset than for other *in vivo* post-crush injury timepoints. This indicates that the timeframes of nerve degeneration during *ex vivo* and *in vivo* degeneration are partially similar, with no differences between sexes. Interestingly, old nerves showed a considerably lower correlation coefficient with the 7dpc *in vivo* datasets than young nerves, potentially indicating an altered degeneration rate in old nerves *ex vivo*. In addition, the proteome changes in old, intact nerves were positively correlated with the respective *ex vivo* degeneration-induced changes for each sex, but in old, male, intact nerves, the proteome changes were slightly stronger positively correlated with *in vivo* degeneration datasets than for old, female intact nerves, suggesting that ageing leads to changes that are partially similar to injury-induced changes, while female nerves appear to be more “protected” from this phenotype (**Figure 2C, Supplementary table 1**).

**Fig. 2.**
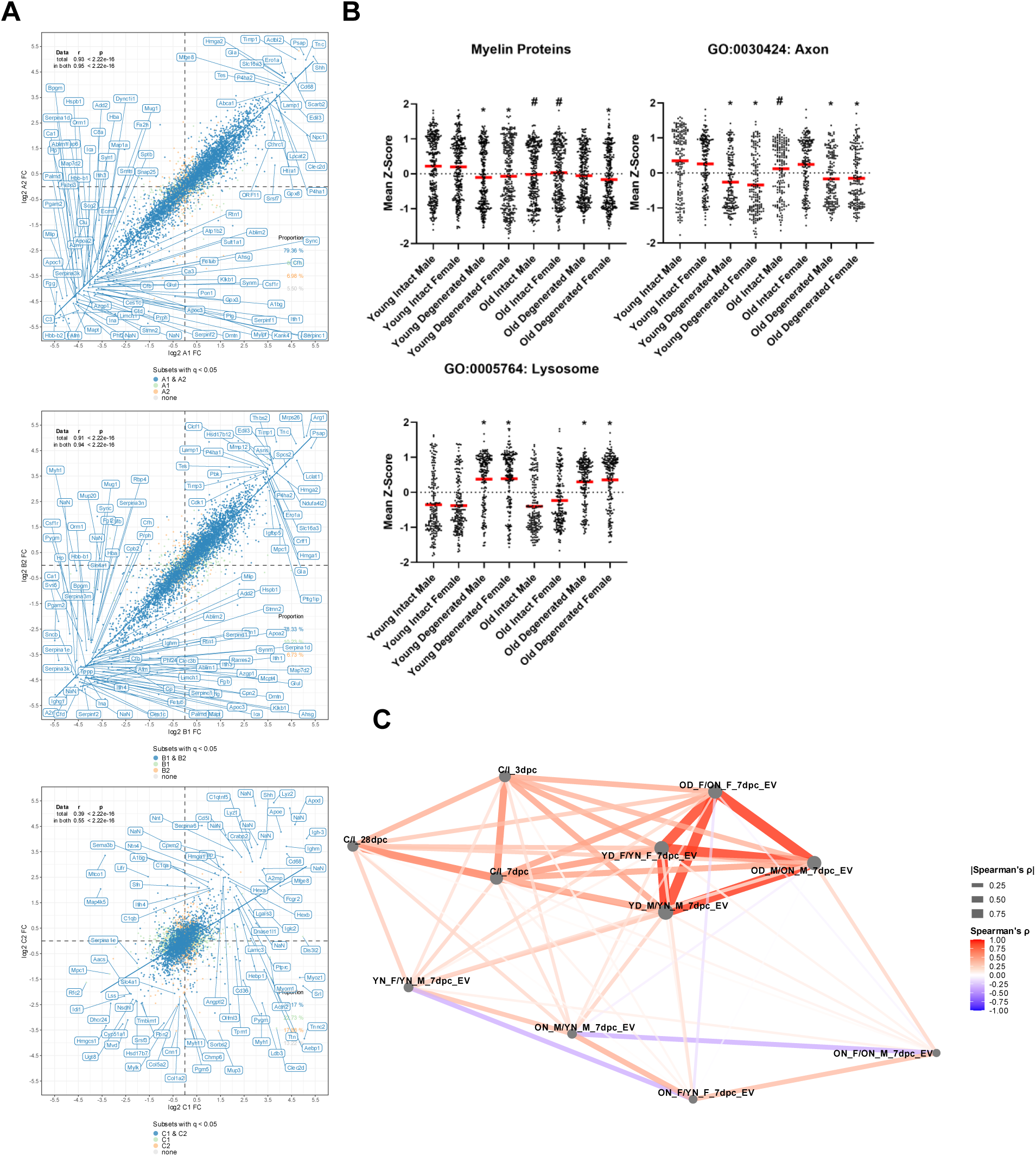
Analysis of the ex vivo degenerative response and correlation to in vivo PNS degeneration. **A** Correlation plot of the log_2_ FCs of the A1 and A2 comparison (upper plot) and B1 and B2 comparison (middle plot) and C1 and C2 comparison (lower plot). In the upper left corner, the correlation statistics for total as well as significantly changed proteins are depicted. In the lower right corner, the percentage of in both comparisons significantly regulated proteins as well as in either male or female mice uniquely regulated proteins is shown. Strongly changed DEPs are labelled with their respective name; Log2FC = log2 fold change; A1 = Young degenerated male / Young intact male, A2 = Young degenerated female / Young intact female, B1 = Old degenerated male / Old intact male, B2 = Old degenerated female / Old intact female, C1 = Old intact male / Young intact male, C2 = Old intact female / Young intact female. **B** Mean z-scores (across all biological replicates from each group) of either all myelin proteins as reported in Helbing et al., 2023 (12) and all axonal and lysosomal proteins that are annotated to the respective GO-terms and were detected in this dataset. Kruskal-Wallis tests for myelin, lysosomal and axonal proteins were performed, each with p < 0.05. P-values from post-hoc tests were FDR-corrected, * indicates p < 0.05 for the comparisons degenerated/intact for each age and sex group, # indicates p < 0.05 for the comparisons old intact/young intact for each sex group. Shown are the mean (red lines). **C** Correlation network of the Log_2_ fold changes of all full proteome datasets for all comparisons during *ex vivo* sciatic nerve degeneration, as well as full proteomic datasets after in vivo degeneration, at different timepoints (3, 7 and 28dpc), as previously reported. Spearman’s ρ is depicted on a colour scale as well as **|** Spearman’s ρ **|** is shown as a function of the line thickness. All correlation coefficients are provided in **supplementary table 1**.

### Enrichment analyses demonstrate the regulation of inflammatory and metabolic proteins following *ex vivo* nerve degeneration

Upon peripheral nerve injury *in vivo*, inflammation and metabolic adaptation are induced, leading to coordinated degeneration and subsequent regeneration (13). To determine the changes that occur during nerve degeneration *ex vivo*, we performed gene ontology (GO) enrichment analyses for biological processes (BP) (**Figure 3A**) and cellular components (CC) (**Figure S2**), as well as KEGG pathway enrichment analyses (**Figure 3B, C**) for all modules and comparisons. As expected, similar enrichment profiles were observed for both the upregulated and downregulated proteins associated with degeneration, independent of age and sex. The lists and module of upregulated proteins primarily contained biological processes associated with RNA and RNA metabolism, as well as metabolism in general, splicing, and vesicle-mediated transport. Specifically, the module of upregulated proteins contained biological processes related to telomere maintenance and mitosis, as well as processes associated with wound healing and repair (**Figure 3A**). Concomitantly, GO terms for cellular components showed enrichment for mitochondrial, vesicular, ribosomal, spliceosomal, DNA/RNA-associated, and proteasomal proteins in the lists and module of proteins upregulated in degeneration (**Figure S2**). Although no biological process GO terms were enriched in the lists of sex-specific upregulated proteins, GO term analysis of cellular components showed an enrichment of mitochondrial and vesicle proteins in young male nerves that were upregulated after *ex vivo* degeneration (**Figure S2**). The lists of downregulated proteins and the module after *ex vivo* degeneration showed an enrichment of GO terms related to biological processes such as inflammation, inflammatory responses and signalling, cell adhesion and migration, immunity and antigen presentation, cytoskeleton organisation, axonogenesis, catabolic metabolism, and vesicle transport and localisation. More GO terms related to lipid metabolism were enriched in proteins that were downregulated in old nerves after *ex vivo* degeneration (**Figure 3A**). Concomitantly, GO terms for cellular components displayed an enrichment for membrane, myelin, lipoprotein particles, axonal and synaptic, as well as cytoskeletal proteins (**Figure S2**). Some GO BP terms were also enriched in the sex-specific downregulated proteins. In young female nerves after *ex vivo* degeneration, for example, biological processes associated with immunity and antigen processing and presentation were specifically enriched in the list of downregulated proteins (**Figure 3A**). Cellular component GO terms primarily showed an enrichment of cytoskeleton and granule proteins in this list of downregulated proteins, in addition to those in the list of downregulated proteins in old male nerves (**Figure S2**). Lastly, although no specific biological processes were enriched in the module unaffected by degeneration, Gene Ontology (GO) Cellular Component (CC) enrichment analysis showed that these proteins mainly belonged to extracellular proteins, the plasma membrane, and G-protein complexes (**Figure S2**). Interestingly, the biological processes enriched in lists of proteins upregulated by ageing, independent of sex, partially resembled the enrichments in the module with proteins downregulated by degeneration with regard to strong induction of inflammatory, defence response, leukocyte activation and other immune-related pathways, as previously reported (13) (**Figure 3A**). In both sexes, metabolic processes related to steroid biosynthesis were enriched in the list of proteins downregulated with ageing (**Figure 3A**). Furthermore, several GO terms associated with inflammatory and wound responses were enriched in old male mice (**Figure 3A**). KEGG pathway enrichment analysis of metabolic (**Figure 3B**), immune system and signal transduction pathways (**Figure 3C**) revealed only weak enrichment of specific pathways in the different lists, apart from the downregulated protein module. Here, several metabolic pathways were enriched, especially inositol phosphate metabolism and valine, leucine and isoleucine degradation, as well as several immune-related pathways such as complement signalling, phosphatidylinositol signalling, ErbB signalling and cell adhesion molecule interaction. Only Sphingolipid metabolism and glycan degradation were enriched in the with degeneration upregulated protein module (**Figure 3B, C**). KEGG pathway enrichment analysis of ageing-induced changes revealed an enrichment of metabolic pathways, specifically steroid biosynthesis, among proteins downregulated with ageing. Conversely, an enrichment of complement signalling and antigen presentation was observed among proteins upregulated with ageing (13). Interestingly, the KEGG pathways pentose and glucuronate interconversions and folate biosynthesis were enriched in the list of proteins uniquely upregulated in old female nerves (**Figure 3B, C**).

**Fig. 3.**
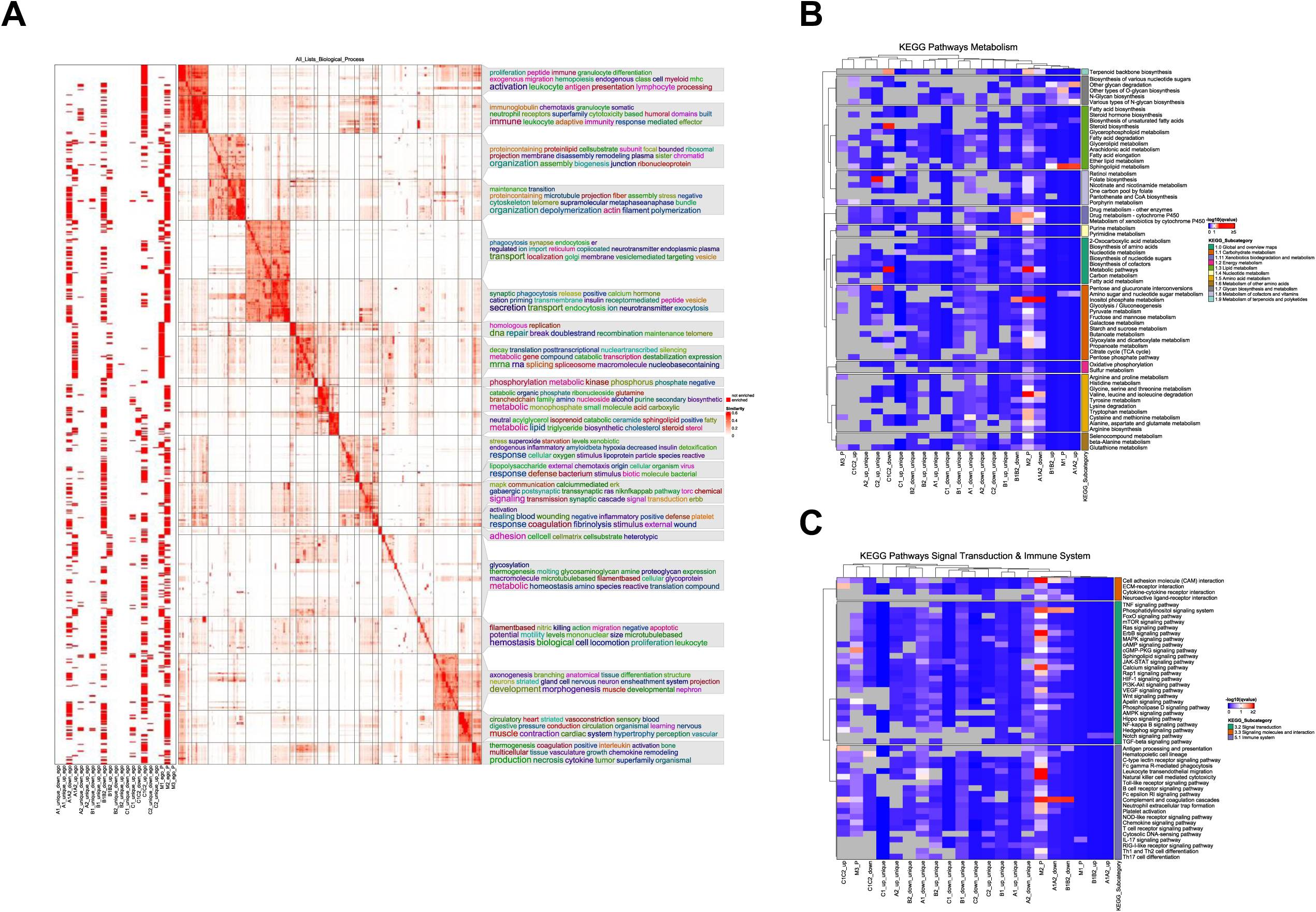
Functional and comparative enrichment analyses of ex vivo sciatic nerve degeneration. **A** Gene ontology enrichment analysis of biological functions for all co-expression modules as well as all the different comparisons among all experimental groups. Semantically similar GO-terms are clustered together (the similarity matrix with specific clusters is depicted in the middle). A binary heatmap on the left shows if a GO-term was significantly enriched in the corresponding gene/protein list (columns). On the right word clouds are annotated to the semantic GO-term clusters with keywords that are overrepresented from all GO-terms located in this cluster, i.e. that serve as representative for the biological functions depicted in this cluster. **B, C** KEGG pathway enrichment heatmaps for all metabolism-related pathways (B) or immune system and signal transduction related pathways (C). In the columns are all gene/protein/module lists and in the rows are all pathways, split by KEGG subcategory. The q-values from the pathway enrichment are shown on a colour scale and gene/protein/module lists that show more similar pathway enrichment profiles cluster together.

## Discussion

In this study, we performed proteomic analyses of sciatic nerves from young and old/geriatric mice and characterized both shared and sex-specific proteomic changes during *ex vivo* nerve degeneration. Overall, degeneration induced extensive proteome remodeling that affected the vast majority of detected proteins, largely independently of age and sex. This finding suggests that the core molecular programme underlying Wallerian degeneration is highly conserved and largely overrides the sex-dependent differences observed in intact nerves (**Figure 1B, Figure S1**). Previous studies have suggested that differences between male and female nerves may arise from sex hormones, such as 17β-estradiol, which can influence Schwann cell differentiation (16). However, another study showed that female mice retain faster nerve regeneration even after ovariectomy (15), indicating that factors beyond ovarian hormones contribute to this phenotype. One possible explanation is sex-specific differences in systemic immune regulation (34, 35): Eliminating all cells, including immune cells, in healthy female mice, ovariectomised female mice and male mice better explained the sex differences in nerve regeneration (15). In either case, the converging responses observed between male and female nerves, and between young and old nerves, can be explained by the absence of sex-specific environmental factors, such as hormone levels and systemic extracellular and serum profiles of soluble factors, in the cell culture dish (35). This is particularly important because systemic ‘inflammageing’ affects peripheral nerve degeneration and regeneration, as we have previously demonstrated (36). Age-related regenerative deficits are thought to arise predominantly from changes in the nerve microenvironment rather than intrinsic neuronal dysfunction. Supporting this concept, heterochronic nerve transplantation studies demonstrated that young mice receiving aged nerve grafts exhibited slower regeneration than recipients of young grafts (5). Consistent with these findings, we observed reduced *ex vivo* myelin degradation in aged nerves, highlighting the diminished capacity of aged Schwann cells and resident macrophages to clear myelin debris (Figure 2B), a defect previously reported in isolated aged Schwann cells (5). Furthermore, Sonic hedgehog (*Shh*) a key regulator of Schwann cell repair phenotype that is essential for efficient degeneration and regeneration (37–39), was among the most strongly induced proteins following degeneration. However, its induction was markedly attenuated in aged nerves. compared to young nerves. Interestingly, intact aged female nerves exhibited higher basal *Shh* abundance than aged male nerves (**Figure 2A** and **Supplementary file 1**), raising the possibility that female Schwann cells maintain a more repair-competent phenotype during ageing. Although speculative, this may contribute to the slower functional decline and enhanced regenerative capacity reported in females (15). The delayed degeneration observed in aged nerves (independent of sex) was further reflected by their weaker correlation with the proteomic profiles of *in vivo* nerve degeneration at 7 days post-crush (dpc), compared with young nerves (**Figure 2C**). *Ex vivo* degeneration remained positively correlated with *in vivo* degeneration across all ages and both sexes, demonstrating that the *ex vivo* model captures many of the intrinsic molecular features of Wallerian degeneration (**Figure 2C**)(8, 40).

Despite these similarities, important differences between *ex vivo* and *in vivo* degeneration were evident, particularly in inflammatory and metabolic pathways. Whereas complement activation and type I interferon signalling are strongly induced during *in vivo* injury and ageing(13)(**Figure 2A, 3C**), these pathways were consistently downregulated during *ex vivo* degeneration ((**Figures 3A** and **3C**, **Supplementary file 2**). This finding suggests that the inflammatory response following peripheral nerve injury depends largely on systemic or blood-derived signals rather than nerve-intrinsic mechanisms alone. Consequently, the *ex vivo* model isolates intrinsic degeneration programmes while highlighting the critical contribution of systemic inflammageing to age-related regenerative decline (36). In contrast, the suppression of metabolic pathways closely resembled that observed during *in vivo* degeneration at 7 dpc (13, 41), indicating that these metabolic adaptations are largely nerve intrinsic. Although most ageing-associated proteomic alterations agreed with our previous findings (13), the present study additionally revealed distinct sex-specific molecular signatures. Aged male nerves exhibited enrichment of inflammatory pathways (**Figure 2A**), consistent with reports of greater systemic inflammageing in males (14). In contrast, aged female nerves showed selective enrichment of the KEGG pathways “Pentose and glucuronate interconversions” and “Folate biosynthesis” (Figure 3B). Enhanced pentose and glucuronate metabolism may improve metabolic flexibility during ageing, as increased activity of this pathway has previously been associated with healthy ageing and rejuvenation, whereas reduced activity has been linked to age-related microbiome alterations (42). The enrichment of folate biosynthesis is particularly intriguing. Folate levels decline with age, and deficiency is common in older adults (43). Reduced folate availability has been associated with central nervous system ageing, depression, cognitive impairment, accelerated epigenetic ageing and peripheral neuropathy (44–46). Moreover, folate supplementation has shown neuroprotective effects in mouse models of diabetic neuropathy (47, 48), and promotes Schwann cell repair programmes following peripheral nerve injury (49). Taken together, these observations raise the possibility that enhanced folate biosynthesis contributes to the greater resilience of aged female nerves (**Figure 3B**) and may partly explain the lower prevalence of age-associated peripheral neuropathy and superior regenerative capacity reported in females (15, 50). Our findings identify folate biosynthesis as a potential molecular pathway associated with healthier peripheral nerve ageing in females. Consequently, targeting folate metabolism may represent a novel therapeutic approach to support healthy peripheral nerve ageing. Nevertheless, this hypothesis remains to be tested experimentally, and future studies are needed to establish whether modulation of folate metabolism can causally improve peripheral nerve regeneration and function during ageing.

### Limitations

Our *ex vivo* degeneration model provides a valuable platform for investigating nerve-intrinsic molecular responses while minimizing systemic influences. However, this is also its principal limitation. Isolated nerves were cultured in standard medium (DMEM supplemented with 5% FBS), which does not reproduce the complex systemic environment present *in vivo*. Consequently, circulating factors known to change during ageing, including inflammatory cytokines (14, 51), as well as sex-specific hormonal differences, were not represented in this model. These missing systemic signals likely contribute to the differences between *ex vivo* and *in vivo* degeneration observed in the present study. Future experiments should address this by supplementing the culture medium with serum from young or aged, as well as male or female, mice to determine how systemic factors influence age- and sex-dependent nerve degeneration.

### Conclusions

To the best of our knowledge, this study provides a comprehensive proteomic characterization of sex-dependent molecular changes during peripheral nerve ageing and *ex vivo* nerve degeneration. Although degeneration activated highly conserved molecular programmes across all experimental groups, subtle but distinct age- and sex-dependent differences were identified. Ageing was associated with enhanced inflammatory signaling in males, whereas female nerves exhibited enrichment of metabolic pathways, including folate biosynthesis, suggesting sex-specific mechanisms that may influence nerve resilience and regenerative capacity.

The *ex vivo* degeneration model reproduced the major proteomic features of *in vivo* Wallerian degeneration, including myelin breakdown, neurodegeneration and activation of cellular clearance pathways. In contrast, inflammatory pathways were suppressed *ex vivo*, indicating that full activation of the inflammatory response following peripheral nerve injury requires systemic signals and/or infiltrating immune cells. Collectively, these findings emphasize the importance of considering biological sex in studies of peripheral nerve ageing and regeneration and provide a valuable resource for investigating molecular mechanisms that promote healthy peripheral nerve ageing.

## Perspectives and Significance

Our study identifies previously unrecognised sex-specific molecular signatures of peripheral nerve ageing and demonstrates that these differences extend to nerve-intrinsic responses during degeneration. The findings suggest that male nerves adopt a more pro-inflammatory ageing phenotype, whereas female nerves preferentially preserve metabolic pathways that may support tissue homeostasis and repair. These molecular differences provide potential mechanistic explanations for the lower prevalence of age-associated peripheral neuropathies and the enhanced regenerative capacity reported in females.

Beyond providing a comprehensive proteomic resource, our work establishes an experimental framework for dissecting nerve-intrinsic and systemic contributions to peripheral nerve ageing. Future studies should validate these findings *in vivo*, investigate the mechanisms underlying the observed sex differences, and determine whether modulation of metabolic pathways, particularly folate metabolism, can enhance peripheral nerve regeneration and promote healthy ageing.

## Acknowledgments

The authors would like to thank Norman Rahnis for excellent technical assistance. Furthermore, the authors would like to thank Madelaine Braune and Claudia Maisch for animal husbandry. Open access funding provided by Projekt DEAL.

## Availability of data and materials

In addition to the data reported in the manuscript and additional files, all the raw proteomic data files are available through the MassIVE platform (**MSV000101693**).

## Supplementary Information

**Supplementary table 1:**
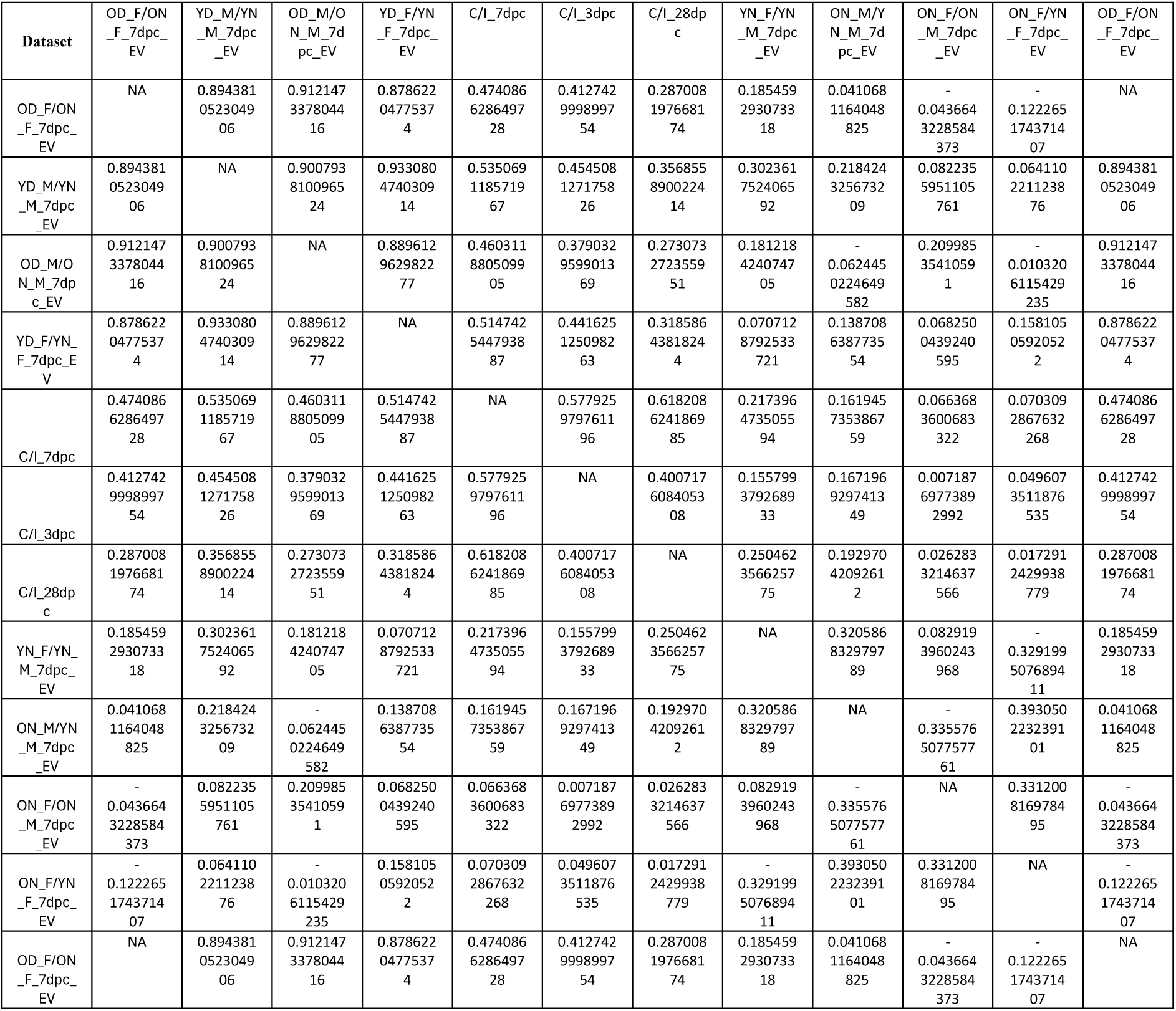
Spearman’s ρ of correlation pairs proteomics datasets.

**Supplementary file 1**: Results from differential expression analysis for all comparisons (candidates as reported by Spectronaut) as well as z-scores of log_2_PG.Quantities across all samples.

**Supplementary file 2**: Results from enrichment analyses for GO BP, GO CC and KEGG pathways.

## Supplementary data figure legends

**Fig. S1.**
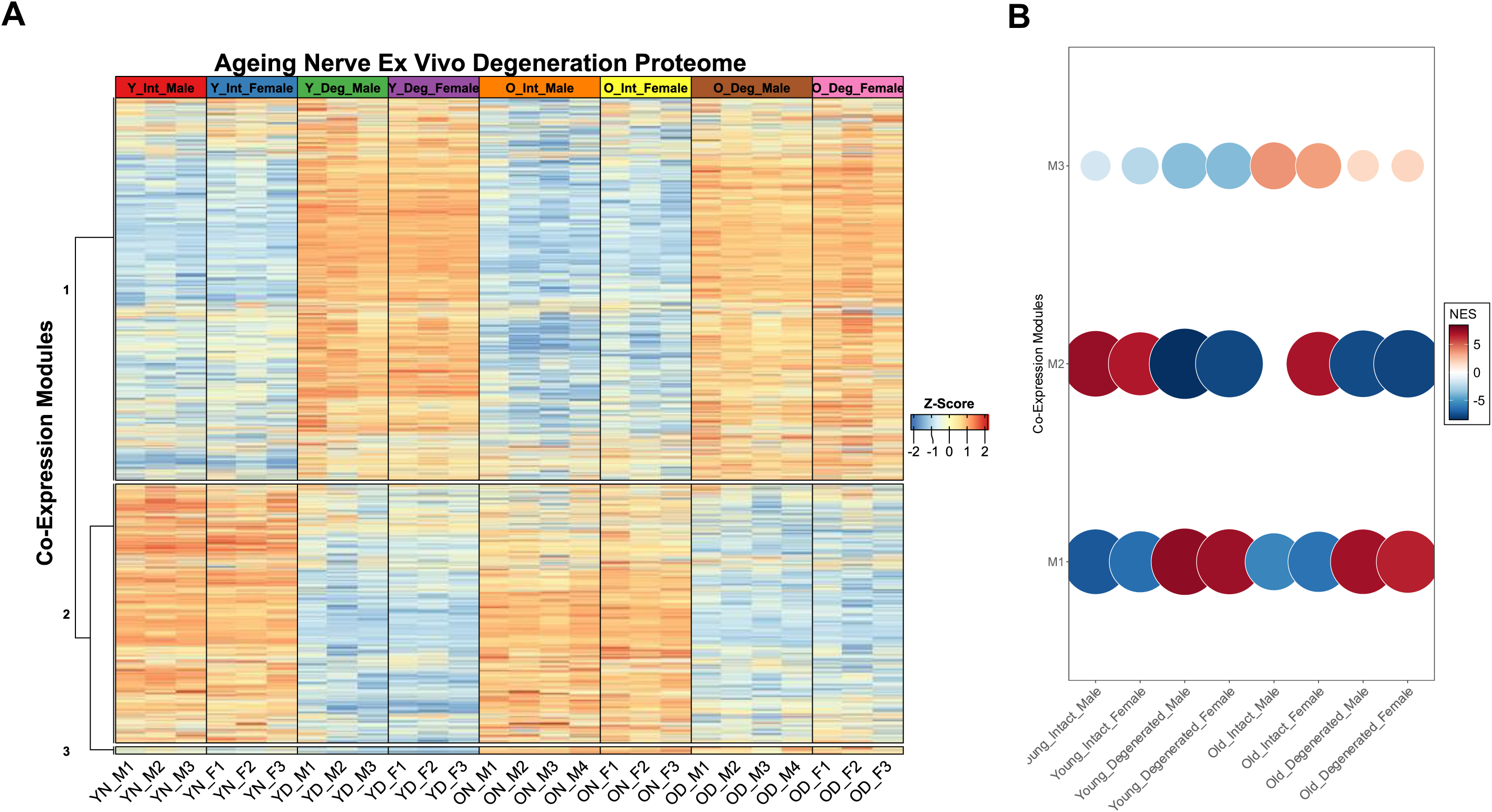
Identification and enrichment of co-expression modules across experimental groups. **A** Identification of different co-expression modules that represent proteins that are co-regulated across different clusters with weighted gene co-expression network analysis (WGCNA). Protein-wise z-score normalized intensities are shown. **B** Numerical enrichment scores for the identified co-expression modules shown in **A** across all different experimental groups; NES could not be computed for old, intact male nerves due to extreme enrichment.

**Fig. S2.**
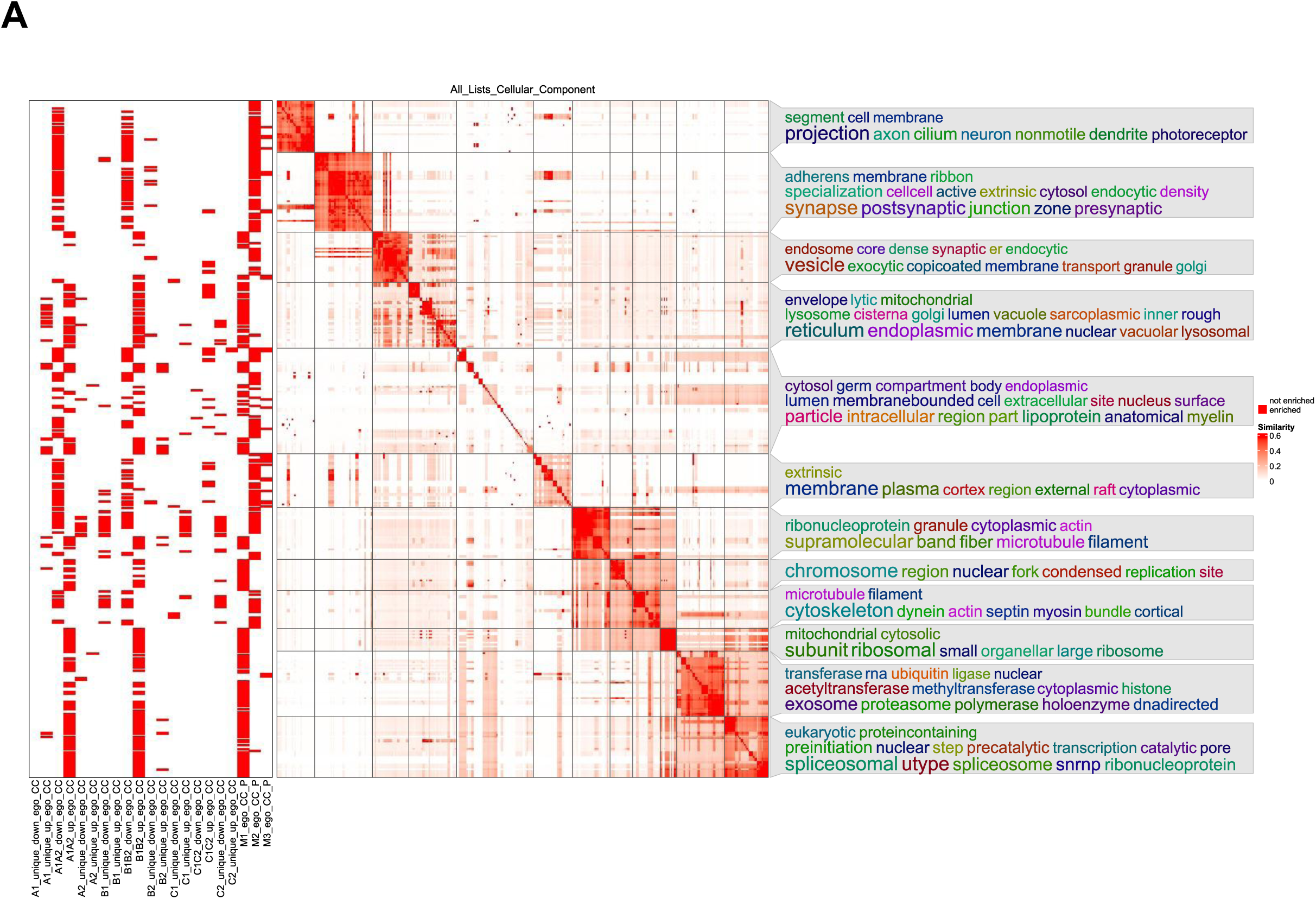
GO-Enrichment of cellular components during ex vivo nerve degeneration. **A** Gene ontology enrichment analysis of cellular component GO-terms for all individual time-specific injury response proteome profiles using semantic similarity analysis, methodically similar to Fig. 3A.

